# Cardiac and behavioural responses to hypoxia and warming in free-swimming gilthead seabream *Sparus aurata*

**DOI:** 10.1101/2021.02.05.429945

**Authors:** Alexandre Mignucci, Jérôme Bourjea, Fabien Forget, Hossein Allal, Gilbert Dutto, Eric Gasset, David J. McKenzie

## Abstract

Cardiac and behavioural responses to hypoxia and warming were investigated in free-swimming gilthead seabream *Sparus aurata* equipped with biologging tags in the peritoneal cavity. After suitable recovery in a holding tank, heart rate (*f*_H_) and the variance of tri-axial body acceleration (VAR_m_) were logged during exposure to stepwise progressive hypoxia or warming, comparing when either swimming in a tank or confined to individual respirometer chambers. When undisturbed under control conditions (normoxia, 21 °C), mean *f*_H_ was significantly lower in tank than respirometers. In progressive hypoxia (100 - 15% oxygen saturation), mean *f*_H_ in the tank was significantly lower than respirometers at oxygen levels until 40%, with significant bradycardia in both holding conditions below this. Mean VAR_m_ was low and invariant in hypoxia. Warming (21 to 31 °C) caused progressive tachycardia with no differences in *f*_H_ between holding conditions. Mean VAR_m_ was, however, significantly higher in the tank during warming, with a positive relationship between VAR_m_ and *f*_H_ across all temperatures. Therefore, spontaneous activity contributed to raising *f*_H_ of fish in the tank during warming. Mean *f*_H_ in respirometers had a highly significant linear relationship with mean rates of oxygen uptake, considering data from hypoxia and warming together. The high *f*_H_ of confined *S. aurata* indicates that static respirometry techniques may bias estimates of metabolic traits in some fish species. Biologging on free-swimming fish revealed novel information about cardiac responses to environmental stressors, which may be closer to responses exhibited by fish in their natural environment.

**SUMMARY STATEMENT:** Implantable biologgers were used to provide the first measurements of cardiac responses to hypoxia and warming in a free-swimming fish, revealing that confinement in respirometer chambers raises heart rate, with consequences for estimates of metabolic rates.

## INTRODUCTION

In fishes the heart is a critical organ for survival, ensuring delivery of oxygen and nutrients to support metabolism, and the removal of metabolic wastes (reviewed by Farrell and Smith, 2017). Cardiac performance is, therefore, considered to be a core determinant of the ability of fishes to survive and thrive in their environment (Farrell and Portner, 2008; Farrell and Smith, 2017; Eliason and Anntilla, 2017; Stecyk, 2017). This includes their ability to tolerate environmental conditions, especially when these become stressful. For instance hypoxia, a reduced availability of dissolved oxygen, is a common stressor in aquatic habitats (Diaz and Rosenberg, 2008) that challenges the ability of the heart to ensure tissue oxygen supply (Randall, 1982; Taylor, 1992). Most fishes are ectotherms, so increases in water temperature have direct thermodynamic effects on their metabolic rate and consequent oxygen demand, which the heart must be able to respond to (Cossins and Bowler, 1987; Schulte, 2011; Rodgers et al., 2016). Investigating how the heart responds to hypoxia and warming is of increasing relevance, because of the hypoxic episodes and summer heatwaves that are occurring in many aquatic ecosystems due to global change (Eliason and Anntilla, 2017; Stecyk, 2017; Costa and Barletta, 2016; Altieri and Diaz, 2018; Stillman, 2019).

The primary cardiac response to hypoxia in fishes is a slowing of heart rate frequency (*f*_H_), known as hypoxic bradycardia (see Taylor, 1992; Farrell, 2007; Stecyk, 2017, for detailed reviews). Although bradycardia is known to be a chemoreflexive response, there is still debate about its actual functional significance for hypoxia tolerance (Farrell, 2007; McKenzie et al., 2009; Joyce et al., 2016; Stecyk, 2017). When warmed, fishes exhibit increased *f*_H_, a tachycardia that may have multiple contributing mechanisms (Eliason and Anntilla 2017). It presumably serves to meet the increased oxygen demands caused by thermal acceleration of metabolism, such that intrinsic capacity to raise *f*_H_ may be a determinant of a species’ upper temperature tolerance (see Eliason and Anntilla, 2017, for a detailed review). Although these cardiac responses to hypoxia and warming have been described in multiple species, this has exclusively been from acute experiments under controlled conditions with animals confined in some way, almost always instrumented with wires connected to a measurement device (Stecyk 2017; Eliason and Anntilla, 2017). It is not known, therefore, whether such cardiac responses to hypoxia and warming would also be observed in fishes free to swim and exhibit behavioural responses to the stressors.

Small biologging tags that record *f*_H_ from the electrocardiogram (ECG) are now available, which can be implanted into fishes to measure their cardiac activity when they are recovered and free-swimming (Prystay et al. 2017, Brijs et al. 2018; Ekström et al. 2018, Bjarnason et al. 2019). Such tags can also log external tri-axial body acceleration (EA), such that it is possible to interpret changes in *f*_H_ against simultaneous measures of spontaneous behaviour (Clark et al. 2010). That is, such tags can measure cardiac responses to hypoxia and warming in fishes swimming freely, and provide insight into how such responses might be affected by spontaneous behavioural reactions to the environmental stressors. This should show cardiac responses that are a more reliable reflection of response patterns by wild animals in their natural environment.

We implanted biologgers into gilthead seabream *Sparus aurata*, to compare cardiac and behavioural responses to progressive hypoxia and warming when the animals were either shoaling in a tank or confined in individual respirometer chambers. We expected that fish would exhibit hypoxic bradycardia and warming tachycardia in both tank and respirometers, but that patterns and thresholds of cardiac responses to the stressors could differ between the holding conditions. In particular, we expected that *f*_H_ of fish would be higher in the tank, due to spontaneous activity, with consequences for cardiac response patterns to the stressors that we were unable to predict with any confidence. Nonetheless, we expected to find an overall direct relationship between activity in the tank, logged as EA, and *f*_H_. We also expected to confirm, using respirometry in the chambers, that *f*_H_ is a predictor of metabolic rate in this species (Hachim et al., 2020).

## MATERIAL AND METHODS

### Ethical approval

Experimental procedures were approved by the ethics committee for animal experimentation n° 036 of the French Ministère de l’Enseignement Superieur, de la Recherche et de l’Innovation, with reference number APAFIS#20294-2019040516446800 v3.

### Animals

Experiments were performed on n = 12 seabream with a mass of approximately 500 g and age of approximately 18 months. The seabream were obtained from the Ferme Marine du Douhet (La Brée les Bains, France) as post-larvae then reared at the Ifremer Aquaculture Research Station in Palavas-les-Flots, in indoor cylindrical tanks (vol 2 m^3^) under seasonal photoperiods, provided with a flow of biofiltered and UV-treated seawater at 21 °C. Fish were fed daily with commercial pellets but fasted for 24 h prior to surgery.

### Surgery

Fish were anesthetized by immersion in 0.1 g l^-1^ benzocaine in aerated seawater, until active ventilation ceased, then weighed and placed on an operating table with their gills irrigated with aerated seawater containing 0.05 g l^-1^ benzocaine. Heart rate loggers (DST milli HRT-ACT, 13 mm × 39.5 mm, 12 g, Star-Oddi, Iceland, www.star-oddi.com) were implanted in the intraperitoneal space, via an incision in the midline below the pectoral fins. Loggers were advanced as close as possible to the pericardium and fixed with sutures (silk suture and non-absorbable monofilament) such that their ECG electrodes lay against the body wall, with the incision then closed with sutures (non-absorbable monofilament). Fish were recovered in a 1 m^3^ cylindrical tank provided with a flow of aerated, biofiltered and UV-treated seawater at 21 °C, with minimal disturbance beyond visual inspection in morning (08:30) and evening (17:00). Fish were not fed during recovery or subsequent experiments. The tank was in a separate room with a natural photoperiod through skylights, shielded behind black plastic curtains with all disturbance kept to an absolute minimum. The timing of recovery is shown in figure 1.

**Fig. 1.**
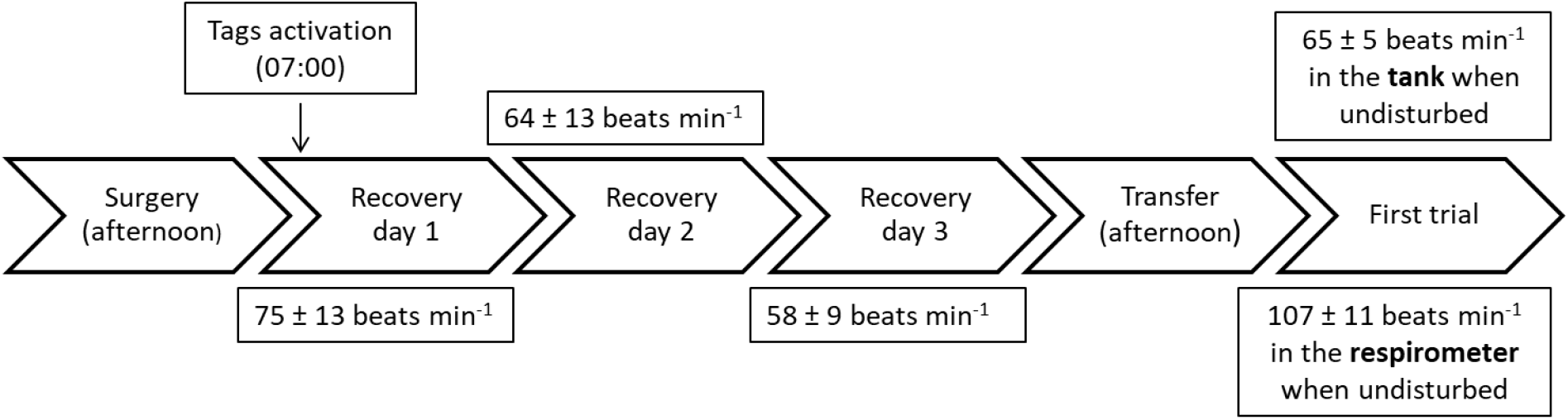
Timeline of fish recovery after surgery, with mean (± S.D.) heart rates (*f*_H_) for specific intervals. Each arrow represents an entire day. The undisturbed value corresponds to the mean *f*_H_ of all fish between 07:00 and 08:00, prior to experiments.

**Fig. 1.**
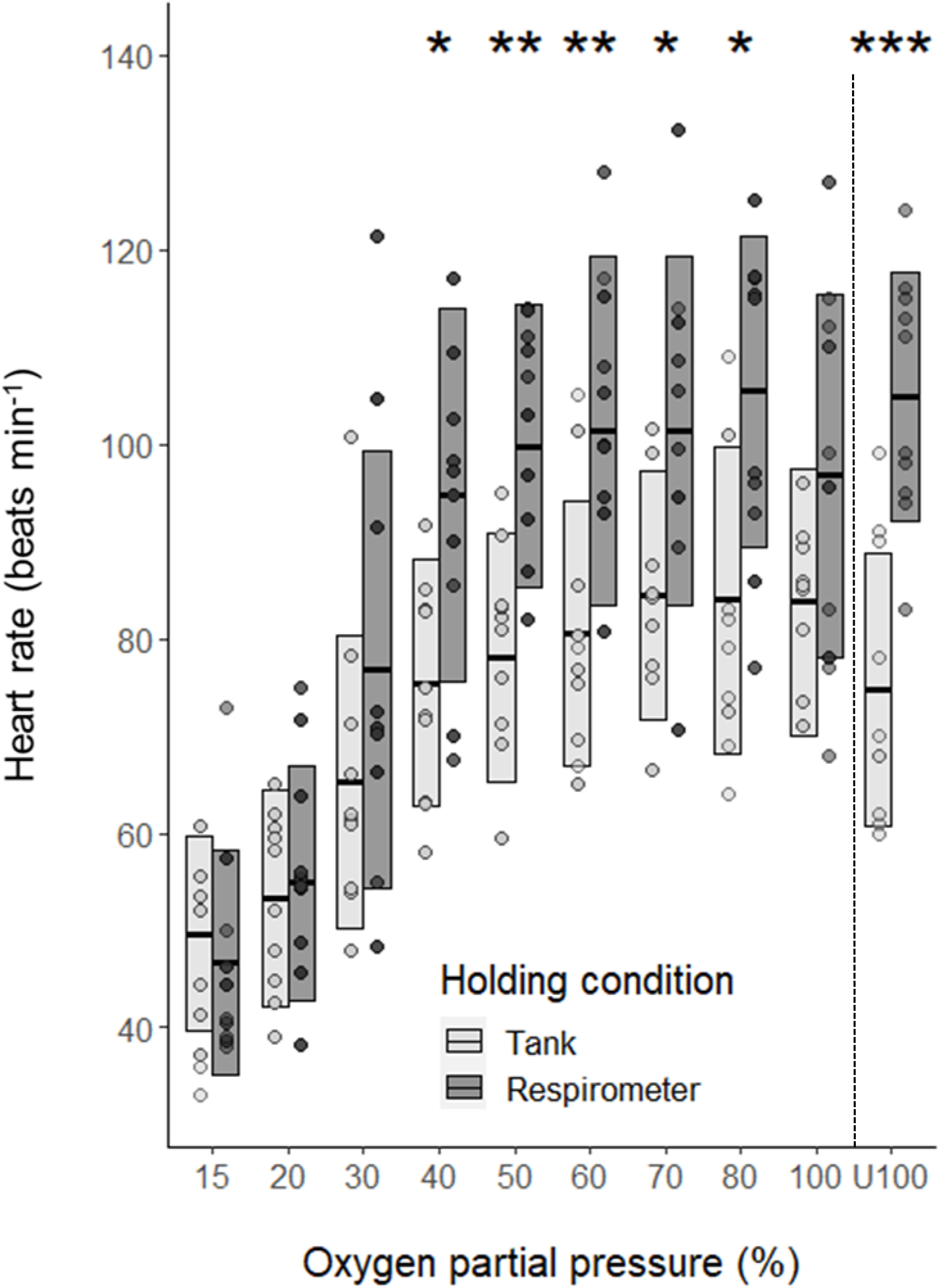
Mean heart rate (*f*_H_) of *S. aurata* during progressive hypoxia while free-swimming in a tank or confined in a chamber. Central bars denote mean values and boxes denote standard deviation, with each point being the mean value of a single fish (n = 10 in all cases). “U100” refers to *f*_H_ of fish when undisturbed in normoxia between 07:00 and 08:00, before hypoxia trials. The vertical dashed line indicates the beginning of the hypoxia trial. Asterisks show oxygen levels where there was a significant difference between tank and chamber; * p <0.05; ** p <0.01.

### Hypoxia and temperature challenges

Fish were instrumented and studied in two groups of six individuals, with each individual tested when swimming freely in the tank and also when confined in a respirometer. On the fourth day after surgery (Fig 1), three of the six fish were netted from the 1 m^3^ tank and placed in individual rectangular respirometer chambers (vol. 9 l) which were submerged in a small raceway in the same room and provided with the same water as the tank. After 24h acclimation to the respirometers (hence at 5 days after surgery for all animals, Fig 1), experiments were conducted over five days. For the first set of six individuals, these had the following sequence: on the first day fish were exposed to a warming challenge, the second to a hypoxia challenge. At the end of the second day, the three fish were exchanged between tank and respirometers and allowed 24 h to recover and acclimate to their new holding condition. On the fourth day they were given a hypoxia challenge, on the fifth, a warming challenge. For the second set of six, the order of exposure was changed to offset any effect of experimental sequence on responses. The order was hypoxia, warming, exchange fish, warming, hypoxia. Black plastic curtains were used to screen tank and respirometers from visual disturbance, fish were observed through small holes in the curtain. Care was taken to reduce all disturbance to a minimum during experiments, experimenters entered at 08:20 to set up the trials then gave fish 30 min to recover from any possible disturbance before commencement.

For progressive hypoxia, oxygen partial pressure in the tank and raceway was decreased simultaneously by bubbling with 100% nitrogen, from 100 % (normoxia) to 80 %, then in steps of 10 % from 80 % to 20 %, then finally 15%. Each step had a duration of 30 min, water oxygen levels were recorded using optical oxygen probes (Firesting sturdy dipping probes, Pyroscience, www.pyroscience.com) and meter (FireSting FSO2-4), with data displayed and recorded in the Pyro Oxygen Logger software, with nitrogen flow and setpoints controlled manually. For acute warming, temperature was raised simultaneously in the tank and respirometers, in steps of 1 °C every 30 min, from 21 to 31 °C, using automated temperature control systems (AquaMedic T controller twin, www.aqua-medic.de) that reached incremental setpoints precisely by controlling activity of a submerged pump, in tank or raceway, that generated a flow of water through heat exchange coils immersed in a reservoir (1 m^3^) of tapwater held at 40 °C. All fish were exposed to these levels of hypoxia and warming; the limits of 15 % saturation in hypoxia and 31 °C in warming were chosen to avoid major stress, which might affect subsequent responses during the 5-day exposure protocol. At the end of the trials fish were euthanized by a lethal dose of benzocaine (1 g l^-1^) to retrieve the logger and data.

Rates of oxygen uptake (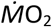, mmol kg^-1^ h^-1^) were measured on the fish in the respirometers, using automated intermittent stopped-flow respirometry (Steffensen, 1989) over a 15 min cycle, providing two measures of 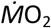 for each 30 min step of hypoxia or warming. Water oxygen concentrations were recorded continuously in each respirometer using Firesting sturdy dipping probes and meter, with data displayed and recorded in Pyro Oxygen Logger. Each fish’s 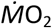 was then calculated considering rate of decline in oxygen concentration in the chamber, chamber volume and the mass of the fish (McKenzie et al., 1995; 2007). Background measurements, on empty chambers, were made prior to placing the fish and at the end of each series. These were always negligible and so no corrections were applied.

### Logger programming and data processing

The loggers were programmed with Mercury software (Star-Oddi) according to the manufacturer’s instructions. For *f*_H_, the ECG data were sampled at 200 Hz for 4 s, once every 5 min from 07:00 to 18:00 during experiments. During recovery from surgery and at night during experiments, *f*_H_ was measured only once every two hours. An ECG trace was saved with each measure of *f*_H_, for visual confirmation of data quality (Fig. S1). For EA, data were collected at 10 Hz for 1 min, once every 5 min from 07:00 to 18:00 during experiments for the first series, but only from 09:00 to 17:00 for the second series, to ensure sufficient battery life. Water temperature, date and time were also recorded with each *f*_H_ and EA measurement.

Heart rate was returned in beats min^-1^, calculated by the manufacturer’s Patternfinder v.1.16.0 software from R-R intervals in the QRST wave of the ECG (Altimiras et al. 1997). Each measure was confirmed by visual inspection of the ECG trace and manual calculation of R-R interval within Patternfinder. The variance of EA (VAR_m_) VAR_m_ was used to identify periods where variation in acceleration indicated bouts of activity or agitation. It was calculated as the variance of the 600 EA measurements per minute, which indicated when the sensor was measuring acceleration above 1 *g*, in units of m*g* (1000 m*g* = 1 *g*), where EA = 0 is equal to 1 *g* and EA = 1000 is equal to 2 *g*. Each measure of *f*_H_ or VAR_m_ was associated with temperature, date and time recorded on the logger. Date and time information on the logger were used to establish the associated oxygen levels for the hypoxia trials, based on the oxygen probe recordings.

### Statistical analysis

Statistics were performed with R version 3.5.3 (R Core Team 2019), with p = 0.05 taken as the fiducial level for statistical significance. A one way analysis of variance ANOVA with repeated measures (stats package, aov function) was used to compare *f*_H_ across recovery days following surgery. Holm-Bonferroni post-hoc tests were used to identify any significant differences among days of recovery. Paired Student t-test was used to compare discrete mean *f*_H_ values between tank and respirometer, namely undisturbed values in the morning prior to trials, plus the maximum and minimum *f*_H_ during trials and the oxygen partial pressure or temperatures at which these occurred. For these tests, normality, homoscedasticity and independence of the residuals were verified visually.

Effects of stressors on *f*_H_ were evaluated and compared between holding conditions by two-way ANOVA with repeated measures (afex package, aov_car function), where one factor was tank versus respirometer and the repeated factor was either oxygen level or temperature. Undisturbed values (21 °C, normoxia) were included in the ANOVA. Normality of the data was verified with a Shapiro-Wilk test. Sphericity of the data was not met, therefore a Greenhouse-Geisser correction was applied. Homoscedasticity of the residuals was verified visually by plotting them as a function of their fitted values. Holm-Bonferroni post-hoc tests were used to identify where significant differences occurred. As a two-way ANOVA with repeated measures does not tolerate missing data, 17 *f*_H_ measures were imputed using either nearest neighbours’ method (kNN function from DMwR package) or linear regression method (see results). These missing and imputed data represented 1.7% of total data. Linear regression models were always significant and calculated *f*_H_ was always plausible.

As VAR_m_ data were not normal, effects of the stressors and comparison between holding conditions were evaluated with a generalized mixed linear model, with fixed factors being holding condition and either oxygen partial pressure or temperature, and individuals as a random effect. Tukey post-hoc tests were used to identify where any significant differences lay.

A linear relationship between VAR_m_ and *f*_H_ was established for all individuals using a generalized mixed linear model, and between *f*_H_ and O2 using a mixed linear model, with individuals as a random effect. Regression slopes between temperature and hypoxia trials were compared using the function lstrends from the lsmeans R package. A linear relationship was also established between *f*_H_ and O2 for each individual, using a linear model. Homoscedasticity and independence of the residuals were verified visually.

## RESULTS

A complete dataset was collected for n = 10 seabream (four from first and six from second series), with a mean (± S.D.) mass of 534 ± 86 g, ranging from 363 to 801 g.

### Routine *f*_H_

Over three days of recovery with absolutely minimal disturbance (Fig 1), overall daily mean (± S.D.) *f*_H_ showed a progressive and significant decrease (p < 0.001) after which some fish were netted and transferred to chambers, to start experiments. For the ensuing five days, fish were necessarily subjected to daily disturbance from presence of experimenters, plus the exposures to hypoxia and warming. Nonetheless, when relatively undisturbed between 07:00 and 08:00 in the morning prior to starting exposures, the mean *f*_H_ of fish was significantly lower in the tank than the respirometers (Tab. 1, Figs 2 and 3).

**Table 1.** Elements of mean (± S.D.) heart rate (*f*_H_, beats min^-1^) of *Sparus aurata* fitted with biologging tags and exposed to hypoxia or warming, comparing when swimming as groups of three in a tank or confined individually in a respirometer chamber. Undisturbed *f*_H_ indicates as measured between 07:00 and 08:00 in normoxia at 21°C, prior to the respective trial. Maximum refers to the mean of the highest, and minimum to mean of the lowest, *f*_H_ observed in each fish in each trial. For hypoxia, PO_2_ at max or min refers to the mean oxygen partial pressure at which maximum or minimum measures occurred, respectively. For warming, T at max or min refers to the mean temperature at which these measures occurred. Asterisks indicate difference between holding conditions for that variable, * < 0.05, *** < 0.001

| Stressor | Holding condition |  |
| --- | --- | --- |
| Hypoxia | Tank | Respirometer |
| Undisturbed | 72 ± 12*** | 105 ± 13 |
| Maximum | 103 ± 10** | 118 ± 13 |
| $\text{PO}_2$ at max | 71 ± 24 | 65 ± 18 |
| Minimum | 37 ± 6.7 | 39 ± 6 |
| $\text{PO}_2$ at min | 17 ± 4 | 19 ± 6 |
| Warming | Tank | Respirometer |
| Undisturbed | 72 ± 15 *** | 103 ± 11 |
| Maximum | 172 ± 22 | 166 ± 17 |
| T at max | 30.3 ± 1.2* | 28.6 ± 2.2 |
| Minimum | 71.5 ± 22 | 81 ± 25 |
| T at min | 21.4 ± 1.8 | 22.8 ± 3.6 |

**Fig. 2.**
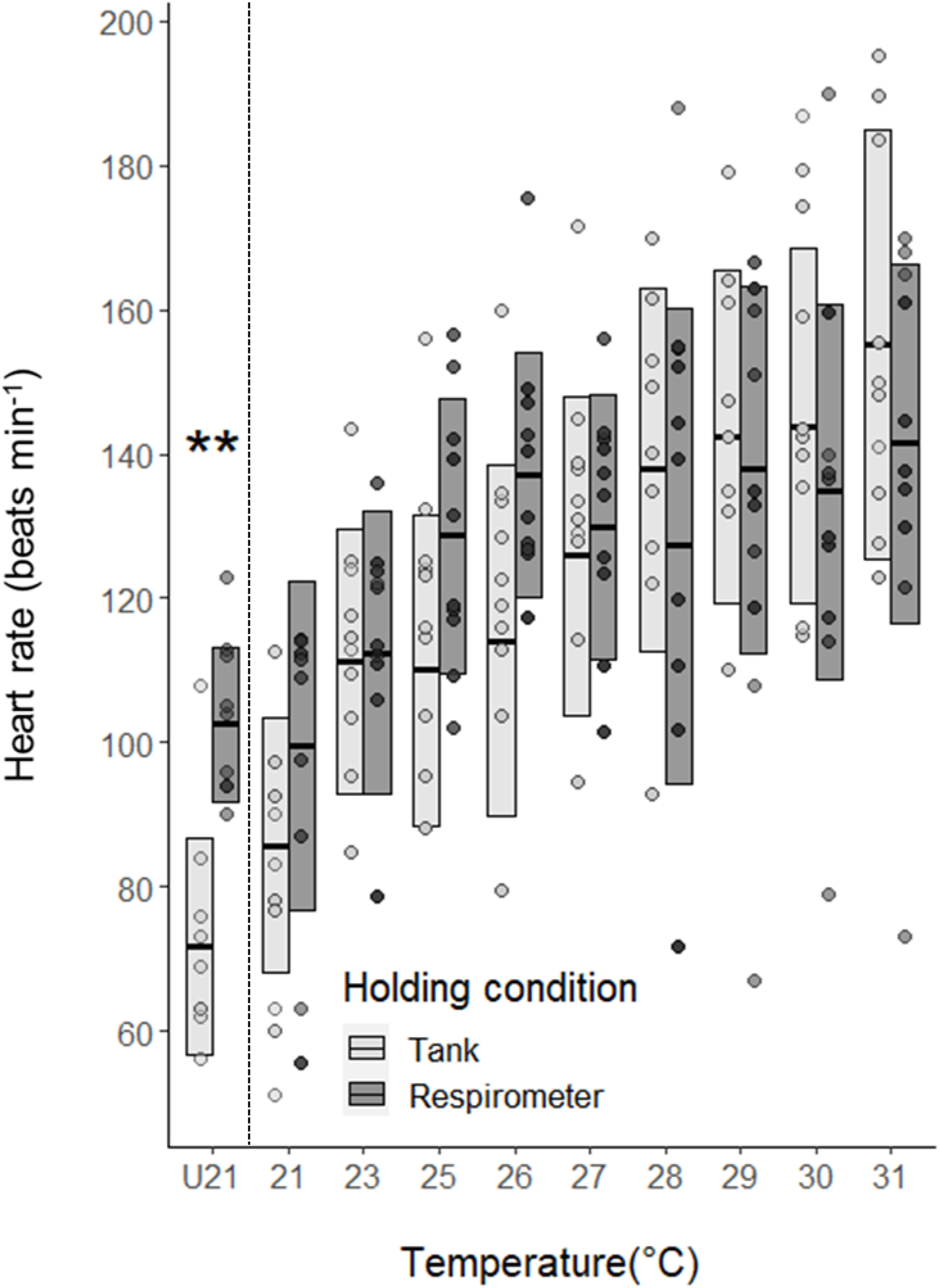
Mean heart rate (*f*_H_) of *S. aurata* during acute warming while free-swimming in a tank or confined in a chamber. Central bars denote mean values and boxes denote standard deviation, with each point being the mean value of a single fish (n = 10 in all cases). “U21” refers to *f*_H_ of fish when undisturbed at 21 °C between 07:00 and 08:00, before temperature trials. The vertical dashed line indicates the beginning of the temperature trial. Asterisks show temperature levels where there was a significant difference between tank and chamber ** p <0.01.

**Fig. 3.**
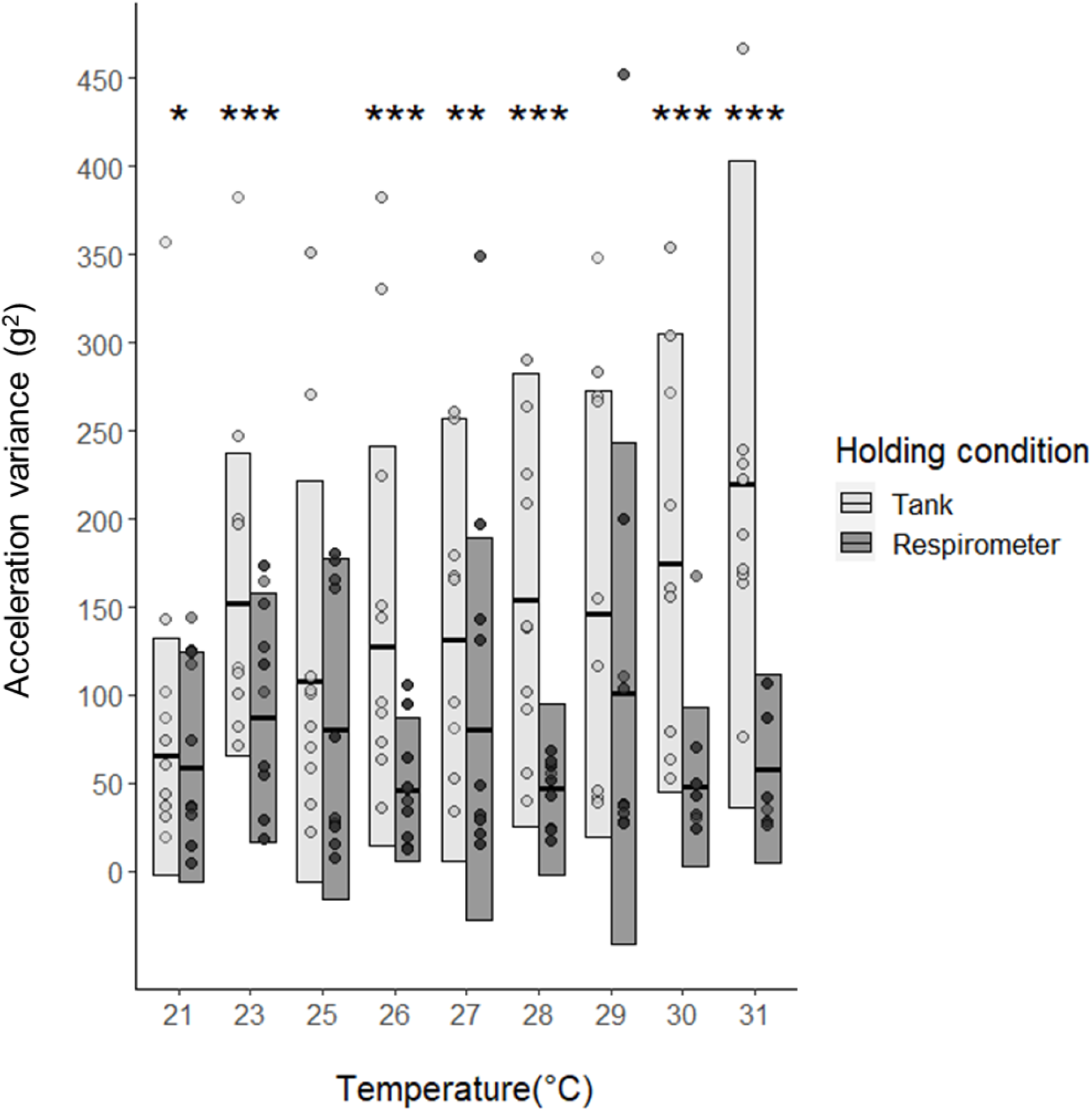
Mean variance of acceleration (VAR_m_) of *S. aurata* during acute warming. Central bars are mean values, each point is the mean value of one fish, and boxes represent standard deviation at each temperature step (n = 10 in all cases). Asterisks show temperature levels where there was a significant difference between tank and chamber * = p <0.05; ** = p <0.01; ***= p <0.001.

### Responses to hypoxia

During progressive hypoxia, *S. aurata* displayed bradycardia in both tank and respirometers (Fig. 2). Within each condition, *f*_H_ did not vary significantly from 100 % down to 40 %, but then *f*_H_ decreased significantly at 30 %, 20 % and 15 %. There was, however, a significant difference in *f*_H_ between tank and respirometer during hypoxia trials (p < 0.01) and a significant interaction between holding condition and oxygen level (p < 0.01). That is, mean *f*_H_ was significantly lower in the tank than in the respirometers at all oxygen steps between 80 % and 40 %. Although undisturbed normoxic *f*_H_ differed significantly (Table 1, Fig 2), this was not true of the measures taken in normoxia as the first step of the exposure trial, notably because of increased individual variation in *f*_H_ in the respirometers (Fig 2). This was presumably because the fish had been disturbed by presence of experimenters. Once bradycardia occurred, namely at 30 %, 20 % and 15 %, there were no significant differences in *f*_H_ between tank and respirometers (Fig. 2). These different patterns of *f*_H_ during hypoxia, between the two holding conditions, were reflected in the fact that mean maximum *f*_H_, whenever this might have occurred during hypoxia trials, was significantly lower (p < 0.01) in the tank than in the respirometers (Table 1). On the other hand, the mean minimum *f*_H_ was similar and occurred at a similar very low oxygen saturation (Table 1). Activity was generally low in hypoxia, with no significant differences in mean VAR_m_ at any level of hypoxia, or between tank and respirometer (Fig. S2). Note that undisturbed values of EA were not collected for all fish but, for four animals, VAR_m_ was low in both conditions, especially so in the tank (data not shown). Visual inspection of the tank revealed that the seabream were moving slowly around the perimeter in hypoxia and tended to stop swimming entirely and rest on the bottom of the tank at hypoxic levels that caused bradycardia.

### Responses to warming

During progressive warming, *S. aurata* displayed tachycardia in both respirometers and tank (Fig. 3). Mean *f*_H_ was statistically similar between holding conditions at all temperatures, despite having been different when undisturbed at 21 °C (Table 1, Fig. 3). Once again, *f*_H_ at the initial 21 °C step of the exposure protocol was variable among individuals, presumably due to mild disturbance. There was, however, a significant interaction between holding condition and temperature (p < 0.01). That is, *f*_H_ increased significantly from 21 up until 27 °C in the tank, but only increased from 21 up until 25 °C in respirometers (Fig 3). Furthermore, the mean temperature at which maximum *f*_H_ occurred was significantly higher in the tank than in the respirometers, being closer to the maximum temperature tested (31 °C) in the tank (Table 1). During the warming trials, VAR_m_ was highly variable in the tank and, at 31 °C, was higher than at all other temperatures (Fig. 4). As would be expected on confined fish, VAR_m_ was relatively low in the respirometers and did not vary significantly with temperature (Fig. 4). As a result, mean VAR_m_ in the tank was significantly higher than in the respirometer at many temperatures (Fig. 4). Visual inspection of the tank showed that the fish were swimming actively around the perimeter at high temperatures, with occasional bursts of speed, especially at 31 °C.

**Fig. 4.**
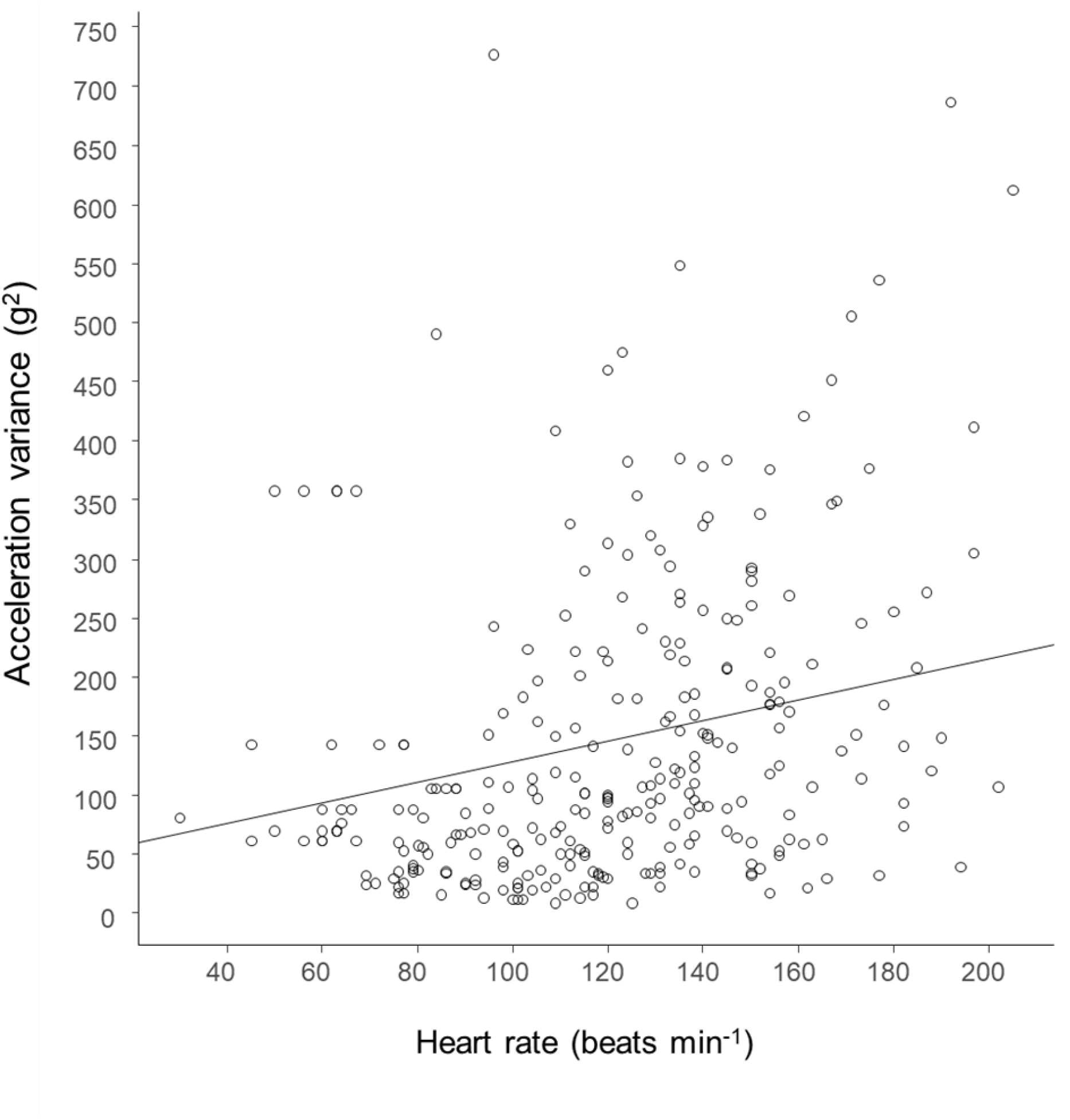
The relationship of variance of acceleration (VAR_m_) to heart rate (*f*_H_) in *S. aurata* during warming trials in the tank. Points are VAR_m_ calculated over 60 s after the corresponding *f*_H_ value for every fish (n = 10 in all cases). The relationship is described by VAR_m_ = *f*_H_ (0.87) + 41.2 (R^2^ = 0.36)

### Relationships between VAR_m_ and *f*_H_, and *f*_H_ and 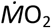

There was a significant linear relationship between VAR_m_ and *f*_H_ in the tank during the warming trials (p<0.001), the only condition where animals showed significant activity (Fig. 5). There was no relation of *f*_H_ to VAR_m_ under any other condition.

**Fig. 5.**
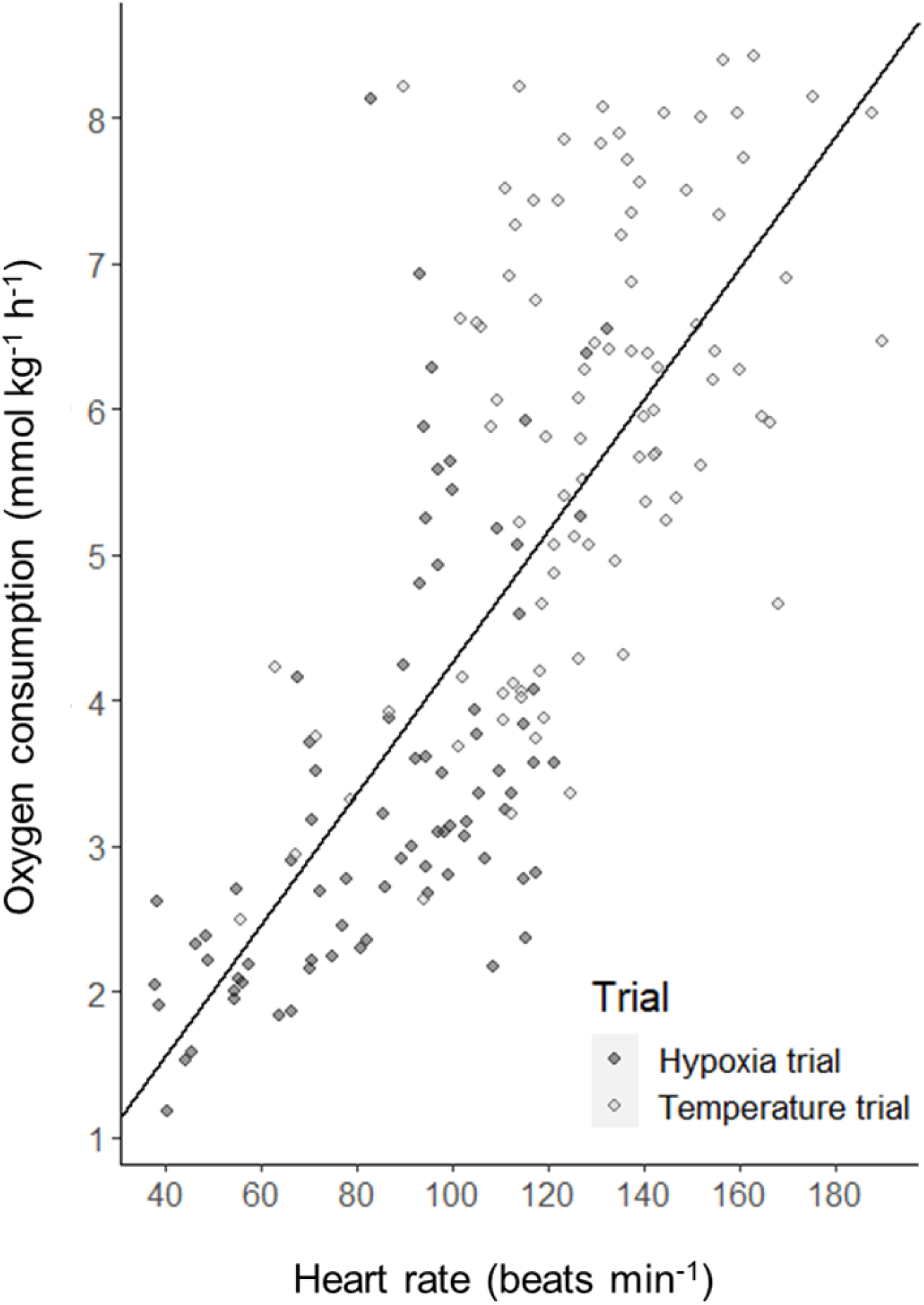
The relationship of oxygen consumption (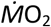) to heart rate (*f*_H_) in *S. aurata* confined in respirometers, during exposure to hypoxia and warming. Each point represents the mean oxygen consumption as a function of mean *f*_H_ for a single fish at a given oxygen partial pressure or temperature (n = 10 in all cases). The relationship is described by 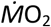 = *f*_H_ (0.045) – 0.23 (conditional R^2^ = 0.76) for all trials combined, represented by the black dashed line.

There was a significant positive linear relationship between *f*_H_ and 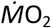 in the respirometers during both hypoxia (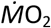 = *f*_H_ (0.034) + 0.4; R^2^ = 0.65; p < 0.001) and warming trials (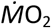 = *f*_H_ (0.02) + 3.37; R^2^ = 0.72; p < 0.001). There was no significant difference between these two slopes, so a single linear relationship between *f*_H_ and 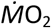 was fitted for all hypoxia and warming values plotted together, which was highly significant (p < 0.001; Fig. 6). Heart rate was also a predictor of metabolic rate for each individual fish (Table S1).

## DISCUSSION

This study is the first report of cardiac responses to hypoxia and warming in a free-swimming fish, although these cardiac loggers have been used on several species (e.g. Norling, 2017; Davidsen et al., 2019; Prystay et al., 2017, 2019; Brijs et al., 2018, 2019; Skeeles et al. 2020). The results supported our expectations, in that fish exhibited hypoxic bradycardia and warming tachycardia in both tank and respirometer. For warming tachycardia, there was clear evidence that increased spontaneous activity levels in the tank could contribute to increased *f*_H_ and affect response pattern. A major unexpected result, however, was that routine *f*_H_ of undisturbed seabream was higher when they were confined compared to free-swimming.

### Routine heart rates and the effects of confinement

The relatively high *f*_H_ measured over a day after surgery presumably indicates an acute stress response, which may have included a release of circulating catecholamines (Reid et al.,1998, Gallo and Civinini 2003) and/or removal of inhibitory cholinergic neural control (Randall 1982, reviewed in Farrell et al. 1984). The progressive decline in *f*_H_ during ensuing recovery presumably indicates an associated decline in stress and recovery of autonomic control (Campbell et al. 2004, 2007; Taylor et al. 2010, reviewed in Sandblom et al. 2011). In free-swimming Atlantic salmon *Salmo salar*, Føre et al. (2020) found that an average of 4 days was required for to recover a stable *f*_H_ after implantation of these loggers. The routine *f*_H_ of the seabream in the third day of recovery, a mean of 59 beats min^-1^ at 21 °C, are amongst the lowest resting values reported for this species (Aissaoui et al. 2000, 2005; Altimiras et al. 1997; Hachim et al., 2020). Comparisons are confounded by differences in size and water temperature, and by our finding that confinement raises *f*_H_ in this species, since all previous studies were on confined fish.

The most obvious explanation for the fact that, when undisturbed, seabream had higher *f*_H_ in the respirometers would be a stress response to confinement, as stress is known to increase heart rate in fishes (Farrell et al. 1991, Sopinka et al. 2016, Rabben and Furevik 1993, Claireaux et al. 1995, Lefrancois et al. 1998, Svendsen et al. 2021). The proximate mechanism for the high *f*_H_ in seabream confined in chambers requires further investigation. This implies, nonetheless, that allowing the seabream to shoal with conspecifics was less stressful than being confined alone. This finding indicates that confinement may introduce bias into studies of physiological responses by fishes to environmental stressors, in a manner that may differ among species. Notably, it could bias measures of metabolic rate by static respirometry, given that *f*_H_ can be a predictor of 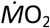, which is the case for *S. aurata* (Hachim et al., 2020). The experiments also revealed how sensitive seabream were to disturbance as, despite taking great care, our simple presence was enough to obscure differences in *f*_H_ between tank and respirometer at trial commencement in normoxia at 21 °C.

### Responses to hypoxia

The data demonstrate that hypoxic bradycardia is observed in free-swimming fish, with a general pattern of response very similar to that observed when confined in a respirometer. Hypoxic bradycardia is a reflex response in teleosts, the sensory arm being chemoreceptor nerve cells in and around the gills that sense oxygen levels in ventilatory water and blood streams and transmit this information to the brainstem. The reflex response occurs via cholinergic fibres in the cardiac branch of the vagus nerve, which slow the heart (reviewed by Taylor 1992; Farrell and Smith 2017; Stecyk, 2017). The functional significance of hypoxic bradycardia is still debated but it may protect function of the cardiac pump, a purely aerobic organ, by conserving contractility and reducing myocardial energy requirements when oxygen supply in the blood is below a critical level (Farrell, 2007; McKenzie et al., 2009; Iversen et al., 2009; Joyce et al., 2016).

It is noteworthy that, although *f*_H_ was significantly lower in the tank compared to the respirometer at oxygen levels above the threshold for hypoxic bradycardia, this threshold did not differ, being between 40 and 30 % in both conditions. The higher *f*_H_ in seabream confined in a respirometer should, presumably, have been accompanied by a higher 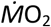 than when swimming freely in the tank, given the direct relationship between these two variables. It might be expected, therefore, that the threshold for bradycardia would be higher in respirometer than in the tank. The fact that the threshold was the same and that, once bradycardia did occur, *f*_H_ was similar between tank and respirometer, requires further investigation.

The low VAR_m_ during progressive hypoxia, and absence of differences between tank and respirometers, presumably indicates that movements in the tank did involve changes in speed, which are necessary to engender variation in acceleration (Tanaka et al. 2001; Hinch et al. 2002; Kawabe et al. 2003). That is, the gentle movements observed during hypoxia in the tank were clearly below the sensitivity of the accelerometer in the tag.

### Responses to warming

Although these tags have been used to study cardiac responses to acute warming in an anaesthetised fish (Skeeles et al. 2020), this is the first report of responses by a fully-recovered free-swimming animal. As for hypoxia, cardiac responses were generally similar between tank and respirometer, with a pronounced tachycardia in both cases. Warming tachycardia in fishes presumably represents a response to increased oxygen demand when metabolism is accelerated by warming, as demonstrated by the linear relationship of *f*_H_ and 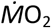 during warming in the respirometer. In terms of the heart itself, this response may reflect both direct effects of temperature on pacemaker function and modulation of autonomic control (see Eliason and Anntilla, 2017, for a detailed review). In the seabream, the maximum *f*_H_ observed during warming, 205 beats min^-1^ at a temperature of 31 °C, was about double the maximum achieved during forced exercise in a swim-tunnel at 16 °C in this species (Hachim et al., 2020). It was also considerably higher than the maximum *f*_H_ typically reported for temperate fishes, of around 140 beats min^-1^ (Casselman et al. 2012, Farrell et al. 2009). These very high *f*_H_ in the seabream were all confirmed by visual inspection of the traces, with clear ECG waveforms (Fig. S1).

It is interesting that, unlike in hypoxia, the accelerometer detected activity in the tank during warming, especially at the higher temperatures. The consistently higher VAR_m_ in the tank relative to the respirometer, for most warming steps, would explain why *f*_H_ did not differ significantly between the holding conditions. That is, the VAR_m_ data confirm that, in free-swimming individuals, some of the tachycardia was due to behavioural responses to warming. The fact that activity contributed to the cardiac response in the tank may explain why the mean temperature for maximum *f*_H_ differed between holding conditions, being higher in the tank and close to the maximum temperature tested, 31 °C. The data revealed a threshold for a behavioural response, with the significant increase in VAR_m_ relative to acclimation temperature, when fish had reached 31 °C. The activity observed in the tank, especially the bursts of speed at high temperatures, may have reflected attempts to escape the conditions, although fish did not become agitated at the same temperature when confined in the respirometer. Such behavioural thresholds may be useful indicators of tolerance that are more sensitive than, for example, loss of equilibrium at a critical thermal maximum (CT_max_). In *S. aurata*, CT_max_ ranges from about 34.3 °C to 36.6 °C, depending on acclimation temperature (Kir et al. 2020, Madeira et al., 2014, 2016)

### Relationships of heart rate to acceleration and metabolic rate

The significant dependence of *f*_H_ on VAR_m_ during warming in the tank is further proof that activity was responsible for raising *f*_H_ of the free-swimming fish. The relationship was, nonetheless, rather noisy with low predictive power. This may be because increases in VAR_m_, especially at high temperatures, reflect agitation and burst swimming movements powered by fast-twitch glycolytic muscle (Webb, 1998). The metabolic costs of such movements are paid during recovery, rather than during the activity itself (Webb, 1998; Kieffer, 2000), so changes in *f*_H_ may have been out of phase with changes in VAR_m_. Although measures of body acceleration have also been used to predict metabolic rate in fishes (Gleiss et al., 2010; Wilson et al., 2013; Wright et al., 2014; Metcalfe et al., 2016; Bouyoucos et al., 2019), it seems unlikely they will ever have the same predictive power as *f*_H_, not least because movement is only one component of metabolic activity in fishes.

The data confirm that heart rate can be used to predict metabolic rate in *S. aurata* (Hachim et al., 2020). The fact that a single linear relationship between mean *f*_H_ and mean 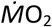 could be described, irrespective of whether data derived from exposure to hypoxia or warming, demonstrates a tight coupling of cardiac pumping activity to metabolic oxygen demand under diverse environmental conditions in this species. The relationships for individual animals were highly significant but their predictive power differed markedly among fish, with variation in *f*_H_ explaining less than 70 % of variation in 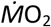 in six of the ten seabream. For this reason, we did not perform the exercise of predicting individual 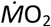 from their *f*_H_ when in the tank. Further research is required to establish the extent to which this variation among individuals is methodological, for example because *f*_H_ and 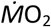 were measured over different time scales, or is physiological. Nonetheless, the results are promising in terms of calibrating the relationship of *f*_H_ to 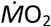 using respirometry and then using logged *f*_H_ data to estimate patterns of energy use by free-swimming seabream (Lucas et al., 1993; Clark et al., 2010; Cooke et al., 2016; Treberg et al., 2016). The need to retrieve the tag is still a limitation on performing such studies on fish released into their natural environment (Prystay et al., 2017, 2019).

## Conclusions

The results demonstrate that hypoxic bradycardia and warming tachycardia are observed in fish whether they are free to shoal in a tank or confined in a respirometer. The fact, however, that confining *S. aurata* in a respirometer raised their routine *f*_H_, presumably due to stress, and that *f*_H_ is a predictor of metabolic rate, has clear implications for estimating metabolic traits by static respirometry in some fish species. Tachycardia in free-swimming fish during warming was due, to some degree, to increased spontaneous activity. That is, the combined measures of *f*_H_ and VAR_m_ in free-swimming fish provided novel insight into drivers of cardiac responses to temperature, and revealed a threshold for a behavioural response to warming. Overall, the results demonstrate that biologging of physiological and behavioural responses to hypoxia and warming, in free-swimming fish, can provide more valid and reliable data than on confined fish, and has potential to reveal sensitive sub-lethal thresholds for impacts of these stressors.

## List of symbols and abbreviations

*f*_H_: heart rate frequency
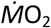: rate of oxygen uptake
EA: external acceleration
VAR_m_: Variance of external acceleration

## ACKNOWLEDGMENTS

The authors are grateful to Marc Vandeputte and Ferme des Douhets for donating the fish. The authors are also grateful Ásgeir Bjarnason of Star-Oddi LTD for technical advice and assistance, and to Germain Salou and Aurélien Leddo of Ifremer Palavas-les-Flots for help in setting up the experiments. The authors also thank Tobias Wang for stimulating discussions about the experiments.

---

**Table S1:**
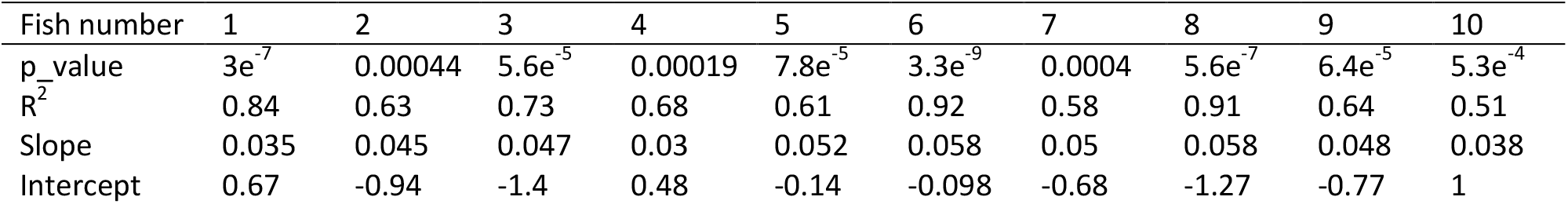
Slope, intercept p-value and R^2^ associated to the linear regression between *f*_H_ and 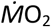 for each fish.

| Fish number | 1 | 2 | 3 | 4 | 5 | 6 | 7 | 8 | 9 | 10 |
| --- | --- | --- | --- | --- | --- | --- | --- | --- | --- | --- |
| p_value | $3e^{-7}$ | 0.00044 | $5.6e^{-5}$ | 0.00019 | $7.8e^{-5}$ | $3.3e^{-9}$ | 0.0004 | $5.6e^{-7}$ | $6.4e^{-5}$ | $5.3e^{-4}$ |
| $R^2$ | 0.84 | 0.63 | 0.73 | 0.68 | 0.61 | 0.92 | 0.58 | 0.91 | 0.64 | 0.51 |
| Slope | 0.035 | 0.045 | 0.047 | 0.03 | 0.052 | 0.058 | 0.05 | 0.058 | 0.048 | 0.038 |
| Intercept | 0.67 | -0.94 | -1.4 | 0.48 | -0.14 | -0.098 | -0.68 | -1.27 | -0.77 | 1 |

**Figure S1:**
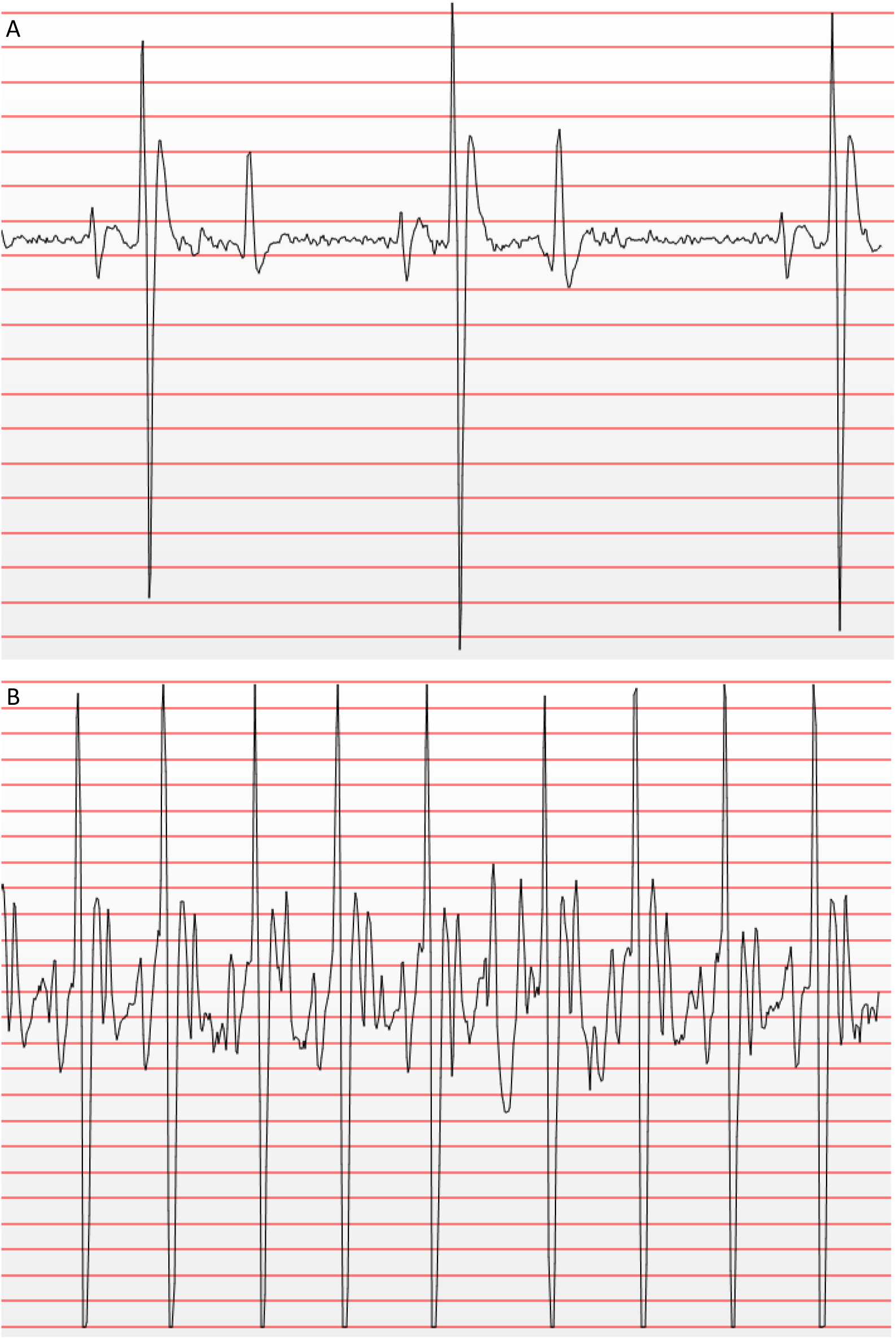
Representative traces of ECG of seabream N°8, mass 500 g, recorded over a 4 s interval by the Star-Oddi 0086 tag, and displayed in PatternFinder^®^ software. Panel A waveform is 51 beats min^-1^ at 21 °C, panel B waveform is 194 beats min^-1^ at 31 °C.

**Figure S2:**
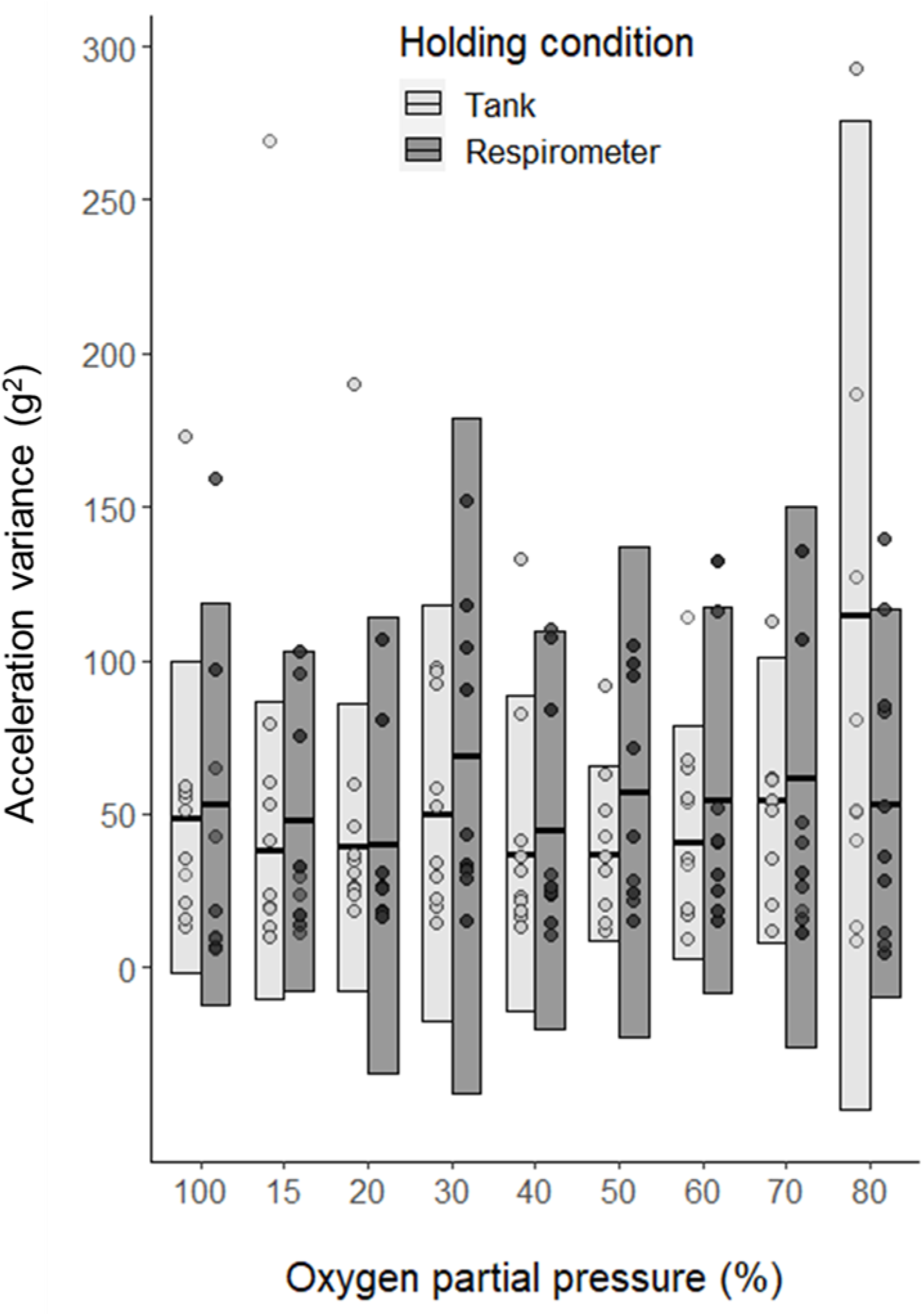
Mean variance of acceleration (VAR_m_) of *S. aurata* during acute hypoxia. Central bars are mean values, each point is the mean value of one fish, and boxes represent standard deviation at each oxygen partial pressure step (n = 10 in all cases).

